# Conformational states during vinculin unlocking differentially regulate focal adhesion properties

**DOI:** 10.1101/176586

**Authors:** Dror S. Chorev, Tova Volberg, Ariel Livne, Miriam Eisenstein, Bruno Martins, Zvi Kam, Brigitte M. Jockusch, Ohad Medalia, Michal Sharon, Benny Geiger

## Abstract

Focal adhesions (FAs) are multi-protein complexes that connect the actin cytoskeleton to the extracellular matrix, via integrin receptors. The growth, stability and adhesive functionality of these structures are tightly regulated by mechanical stress, yet, despite the extensive characterization of the integrin adhesome, the mechanisms underlying FA mechanosensitivity are still poorly understood. One of the key candidates for regulating FA-associated mechanosensing is vinculin, a prominent FA component, which was proposed to possess either closed (“auto-inhibited”) or open (active) conformations. However, a direct demonstration of the nature of conformational transition between the two states is still absent. In this study we combined multiple structural and biological approaches to probe the transition from auto-inhibited to active conformation, and determine its effects on FA structure and dynamics. We further show here that the closed to open transition requires two sequential steps that can differentially regulate FA growth and stability.

## Introduction

Focal adhesions (FA) are subcellular transmembrane structures that anchor the actin cytoskeleton to the underlying extracellular matrix (ECM) via trans-membrane integrin adhesion receptors (Abercrombie and Dunn, 1975; Abercrombie et al., 1971). FA formation is initiated by heterodimeric integrins, which bind to the ECM through their extracellular domains, and are linked to the intracellular actin cytoskeleton via a multitude of associated proteins, commonly referred to as the integrin adhesome (Geiger et al., 2009; Geiger and Zaidel-Bar, 2012; Horton et al., 2016; Humphries et al., 2009; Kuo et al., 2011; Zaidel-Bar and Geiger, 2010). The formation of mature FAs depends on multiple molecular interactions, and is governed by internal and external forces that affect the dynamics of adhesome components and, consequently, the structural integrity of the developing adhesion site (Balaban et al., 2001; Livne et al., 2014; Riveline et al., 2001).

It was shown that after an initial engagement of integrins with the ECM, at nascent adhesions, a dimer of the adaptor protein, talin, binds to the membrane, driving a force-dependent recruitment of multiple adhesome components (Golji and Mofrad, 2014). The mechanism whereby mechanical force activates FA assembly and maturation is rather complex and involves conformational modulation of several FA molecules. Talin, an essential component of FA assembly and of adhesion-mediated signaling, was directly shown to bind to ligand-associated integrins and recruit an increasing number of vinculin molecules upon physical stretching of cells (del Rio et al., 2009; Hu et al., 2016). This is proposed to induce a force-dependent exposure of a large number of vinculin binding sites on talin. With the assembly of FA-associated cytoskeleton, local forces, acting on the nascent adhesions are believed to induce a conformational change within the vinculin molecule, causing its transition from an auto inhibited, closed conformation to an active, open one (Cohen et al., 2005; Cohen et al., 2006). A similar process occurs within cadherin-mediated cell-cell adherens junctions, where vinculin binds to α-catenin to reinforce the adhesion site by recruiting additional proteins (Dufour et al., 2013; Maki et al., 2016; Thomas et al., 2013; Yao et al., 2014). Additional mechanosensitive proteins, like focal adhesion kinase (FAK) and Cas participate in FA regulation, following the force-mediated exposure of their kinase domain, and sequestered phosphorylation sites, respectively (Sawada et al., 2006; Zhou et al., 2015).

Taken together, these studies provided compelling evidence for the key importance of mechanical force in regulating FA formation and fate, yet the molecular basis underlying each of these systems, and the cross-talk between them are still controversial and poorly understood. The activation mechanism of vinculin, is the focus of this work, whereby we investigated its transition from the closed, auto-inhibited state to the open, active conformation.

Vinculin is composed of a globular N-terminal head domain, which consists of 4 α-helical bundles termed D1-D4, and is connected to a C-terminal tail via a flexible hinge (Figure 1A) (Bakolitsa et al., 2004; Borgon et al., 2004). Previous studies have shown that each of these domains can bind to distinct partner proteins (see: http://www.adhesome.org/interactions/VCL.html): The head domain can bind to talin (Fillingham et al., 2005) and α-actinin (Bois et al., 2005), the hinge region interacts with Arp2/3 (DeMali et al., 2002), vinexin (Kioka et al., 1999), ponsin (Mandai et al., 1999) and VASP (Huttelmaier et al., 1998), and the tail domain can bind to actin (Thievessen et al., 2013), paxillin (Subauste et al., 2004) and phosphatidyl inositol bis-phosphate 4,5 (PIP2) (Palmer et al., 2009). Furthermore, the crystal structure of vinculin has revealed that the head domain can interact with the tail, thus forming a stable closed, “auto-inhibited” conformation, which can be disrupted via a high local excess of vinculin binding site of talin (VBS3) (Bakolitsa et al., 2004; Cohen et al., 2005). However, the different vinculin conformers have so far been indirectly inferred from biochemical studies (Chen et al., 2005) and direct characterization of the open, active state is still missing. Nevertheless, the biochemical evidence accumulated so far is in line with the view that interruption of intramolecular bonds between the tail and both D1 and D4 head domains is needed for vinculin activation (Cohen et al., 2005).

**Figure 1.**
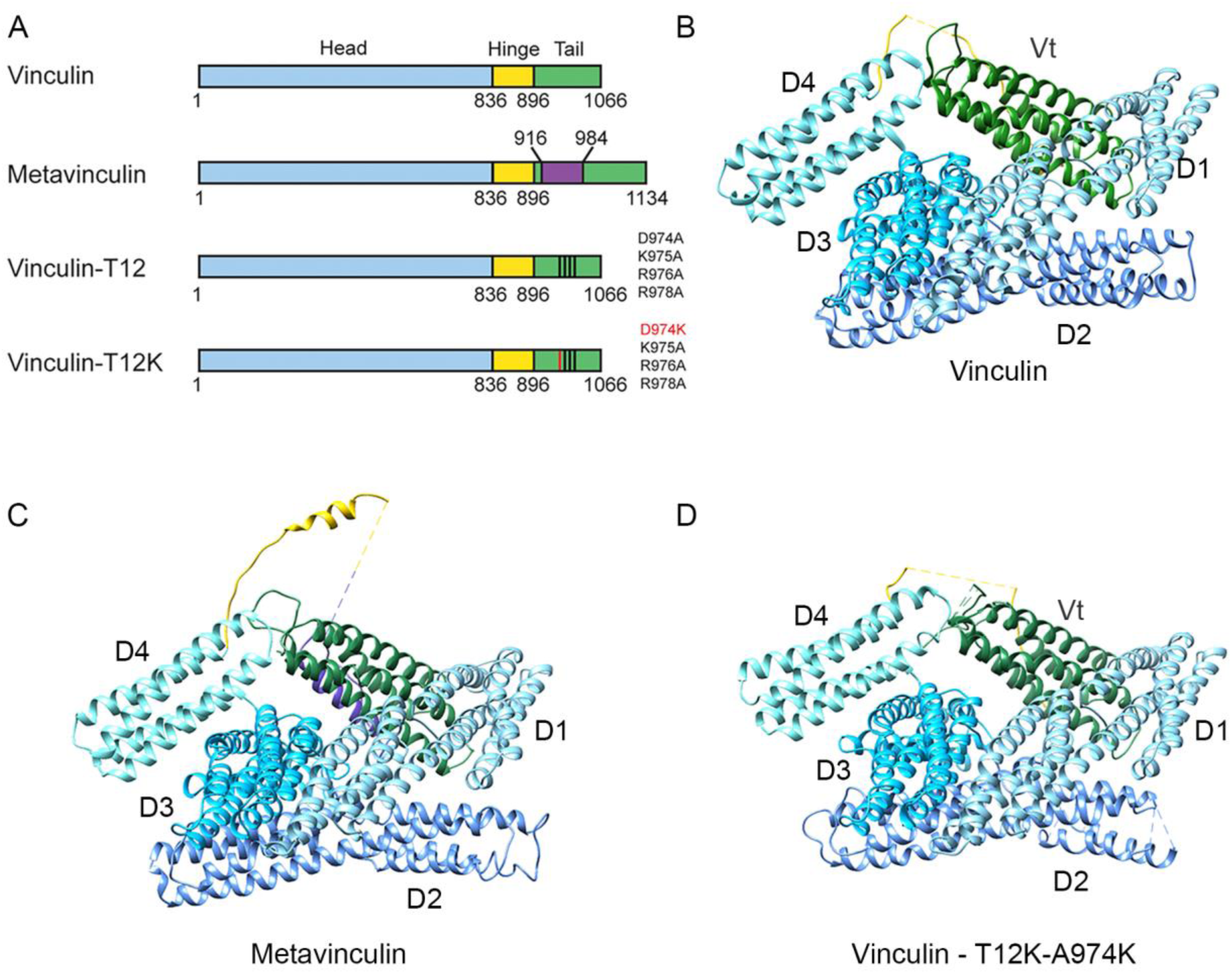
The three dimensional structures of vinculin variants. (A) Schematic illustration of the different vinculin variants, depicting the head (blue), hinge (yellow) and tail (green) domains and the sequence variation between the forms. The 68 amino acid insert of metavinculin is labeled in purple. (B-D) Ribbon depictions of the X-ray structures of vinculin (PDB code 1TR2), metavinculin (Rangarajan et al., 2010) and vinculin-T12-A974K (solved in this study) showing the very similar overall fold. The head, hinge, metavinculin insert and tail demains are colored as in 1A. Missing segments, which were not resolved in the electron density map, are shown as dashed lines.

To investigate the molecular mechanics that mediates vinculin’s transition from the closed to the open state we have compared the energy levels required for opening WT vinculin, to those required for opening a mutant vinculin molecule, termed vinculin-T12. In T12 the head-binding domain on the vinculin tail was mutated (Cohen et al., 2005); see Figure. 1), thereby weakening the head-to-tail interaction affinity by a 100-fold (Cohen et al., 2005). Assays inspecting the kinetics of this mutant as compared to the WT protein, have shown that the lifetime of vinculin-T12 in FAs is three times longer than that of the wild type, implying that vinculin-T12 maintains an open conformation. Hence, binding sites to diverse associated proteins sequestered to the head-tail interface in the closed (“inhibited”) conformer become available for interaction. Further investigation revealed that the presence of vinculin-T12 leads to an extended lifespan of multiple adhesion components in FAs (Carisey et al., 2013; Cohen et al., 2006), as well as increased cell size, surface traction and adhesion strength (Dumbauld et al., 2013).

Given that vinculin and vinculin-T12 were proposed to adopt different conformations, we have chosen to study these two proteins. As an additional tool for studying this process we have created a novel variant of T12, T12-A974K, which contains an additional positive charge that might further destabilize the head-to-tail interactions, and thus, leading to facilitated activation. Beside these two synthetic forms designed to directly probe intramolecular head-to-tail interactions, we also explored the contribution of the hinge region to vinculin activation, by studying the splice isoform metavinculin (mVin), which contains an extra acidic stretch of 68-amino acids placed between the tail and the flexible hinge (Gimona et al., 1987; Gimona et al., 1988; Siliciano and Craig, 1986) (Figure 1A). mVin is present only in smooth, cardiac and skeletal muscle cells, where the cytoskeletal force regime might differ from the one present in non-muscle cells. Previous work suggested that the extra segment of mVin interferes with the auto-inhibitory head-tail interaction and the binding to other classical partners, as well as the ability to specifically bind lipids (Witt et al., 2004). Overexpression of mVin also altered FA structure and the mode of actin bundle organization in cultured cells (Janssen et al., 2012; Rudiger et al., 1998).

In this report, we combined crystallographic studies, native mass-spectrometry (MS), ion-mobility MS (IM-MS) measurements and molecular modeling with cell biological characterization, aiming to understand the mechanism of vinculin activation. Utilizing crystallographic data, we demonstrate that all three variants (T12, T12-A974K and mVin) exist in a closed conformation, similar to that of native vinculin. However, unlike vinculin, T12 and T12-A974K that reside, in solution, predominantly in closed state, mVin coexists in a diverse ensemble of conformations, ranging from the closed state to a range of extended conformations. In addition, collision-induced unfolding experiments coupled with IM-MS measurements show that the opening of all vinculin forms is governed by a two-step unlocking of the head to tail interaction, revealing a distinct “semi-open” conformational state of the molecule. The functional implications of these results are further highlighted by cellular analysis that relates specific steps in the unfolding process to the formation, stability and mechanosensitivity of focal adhesions.

## Results

### A closed head-to-tail conformation is revealed in the crystal structures of vinculin, metavinculin and vinculin-T12-A974K

In an attempt to examine the open conformation of vinculin we have subjected the vinculin-T12-A974K variant, which is expected to comprise the weakest head-tail interaction, to X-ray crystallography (Figure 1D). Surprisingly, the crystal structure we obtained for this variant was of a close conformation, nearly identical to those of native vinculin (Borgon et al., 2004) and metavinculin (Rangarajan et al., 2010) (Figure 1B-D). The structures show that all vinculin variants consist of five helix-bundle domains four of which (D1-D4) form the vinculin head (Vh) that binds through a long, partially unresolved, linker (“hinge”) to the fifth helical bundle (tail domain; Vt). In the closed conformer, Vt appears as a rod whose two ends bind to head domains D1 and D4, via residues conserved in vinculin and metavinculin; a small additional contact is made with D3. The properties of the Vt-D1 and Vt-D4 interfaces differ considerably. The Vt-D1 interface is large (2085 Å^2^, averaged for the two independent polypeptide chains in structure 1TR2), and mostly hydrophobic; yet it is reinforced by hydrogen bonds between positive Vt residues and negative D1 residues. The Vt-D4 interface is considerably smaller, 696 Å^2^ (on the average) and largely electrostatic, positive on the Vt side and negative on the D4 side.

The T12 and T12-A974K mutation sites (residues D974, K975, R976 and R978) map to the Vt-D4 interface and hit hydrogen bonds that stabilize this interface (T974-N773, K975-E772 and R978-E775). The loss of these hydrogen bonds also leads to lengthening of hydrogen bonds between backbone atoms in T12-A974K (Table 1). Hence, the Vt-D4 interaction in the vinculin mutants T12-A974K and T12 is likely to be considerably weaker than in wild type vinculin.

**Table 1:** Hydrogen bond interactions at the Vt-D4 interface as seen in the crystal structures of vinculin, metavinculin and vinculin-T12-A974K (Å)

| vinculin<sup>1</sup> | metavinculin | vinculin-T12-A974K |
| --- | --- | --- |
| T973/O – E775/N<br>2.89<br>2.75 | T1041/O – S774/Oγ 3.26 | T973/O – E775/N 3.65 |
| D974/Oδ – N773/Nδ<br>3.04<br>3.04 | D1042/Oδ – N773/Nδ 3.26 | |
| K975/N – N773/O<br>2.84<br>3.01 | K1043/N – N773/O 3.03 | A975/N – N773/O 3.28 |
| K975/Nζ – E772/O<br>2.35<br>2.29 | K1043/Nζ – E772/O 3.02 | |
| R976/N – N773/Oδ<br>2.94<br>3.03 | R1044/N – N773/Oδ 3.01 | R976/N – N773/Oδ 3.63 |
| R978/Nh – E775/Oε<br>2.62/2.71<sup>2</sup><br>3.00/3.19<sup>2</sup> | R1046/Nh – E775/Oε 2.47 | |

<sup>1</sup>For vinculin two values are listed for the independent polypeptide chains in the asymmetric unit of structure 1TR2
<sup>2</sup>bifurcated hydrogen bond

We further analyzed the Vt-D1 and Vt-D4 interfaces of native vinculin by anchoring spot mapping (Ben-Shimon and Eisenstein, 2010). An anchoring spot consists of a cavity on the surface of a protein (or protein domain) that binds a protruding amino acid side chain of another protein (or domain), and the mapping tool determines the location and binding energy of the protruding amino acid. Anchoring spots with ΔG≤4 Kcal/mol were shown to correspond to very strong experimental hot spots (Ben-Shimon and Eisenstein, 2010). Table 2 lists thirteen strong anchoring spots in the Vt-D1 interface. Seven of them (K944, R945, I997, T1000, T1004, R1008 and E1015) are Vt residues that bind in cavities on the surface of D1, and the other six (P2, F4, E10, E14, E28 and R113) are D1 residues that bind in cavities on the surface of Vt. The Vt-D4 interface includes only three strong anchoring spots, which consist of D4 residues N773 and E775 and Vt residue K975. Anchor residues N773 and E775 form hydrogen bonds to the side chains of D974 and R978, respectively; mutation of these residues to alanine would significantly weaken the Vt-D4 interactions in vinculin-T12 and vinculin-T12-A974K compared to native vinculin. In T12-A974K the additional positive charge augments the positive potential on the Vt side of the Vt-D4 interface. The structure, however, shows that K974 does not form a hydrogen bond with D4; rather, hydrophobic contacts between the CH_2_ groups of K974 and the side chain of N773 are observed. The number of contacts involving K974 exceeds the number of contacts that A974 can make and together with the augmented positive potential it is possible that the Vt-D4 contact in vinculin-T12-A974K is stronger than in vinculin-T12.

**Table 2:** Anchor residues that correspond to calculated low ΔG (in Kcal/mol) anchoring spots*) The ΔG values are averages of anchoring spots calculated for the two independent polypeptide chains in the asymmetric unit of structure 1TR2; an asterisk indicates that an anchoring spot was detected for only one of the chains.

| The Vt-D1 interface |  | The Vt-D4 interface |  |
| --- | --- | --- | --- |
| Residue | $\Delta G^1$ | Residue | $\Delta G$ |
| K944 | -4.0* | N773 | -3.5 |
| R945 | -5.8* | E775 | -3.2* |
| I997 | -3.6 | K975 | -4.1* |
| T1000 | -3.1 |  |  |
| T1004 | -3.3 |  |  |
| R1008 | -4.9* |  |  |
| E1015 | -3.2 |  |  |
| P2 | -3.9 |  |  |
| F4 | -3.8 |  |  |
| E10 | -3.3 |  |  |
| E14 | -3.9 |  |  |
| E28 | -8.8 |  |  |
| R113 | -3.5* |  |  |

\*) The $\Delta G$ values are averages of anchoring spots calculated for the two independent polypeptide chains in the asymmetric unit of structure 1TR2; an asterisk indicates that an anchoring spot was detected for only one of the chains.

### Metavinculin and vinculin-T12-A974K occupy a larger conformational range compared to vinculin and vinculin-T12

Considering that crystal structures, cryogenically frozen before analysis, often reflect only a subset of possible protein conformers (Fraser et al., 2011), we were wondering whether the closed conformation displayed by all vinculin variants, is indeed the only occurring stable state of these proteins in solution. To address this issue we applied the ion mobility-mass spectrometry (IM-MS) approach, which enables the characterization of the ensemble of conformational states of a system in aqueous conditions (Bohrer et al., 2008; Wyttenbach et al., 2014). In this method, the time it takes a protein to transverse a weak electrical gradient in a gas-filled chamber is measured. The drift time is proportional to the number of gas molecules the protein collides with along its path, so that proteins of similar mass, but with less compact conformation will traverse the IM chamber more slowly. As the number of collisions is proportional to the surface area of the protein, a rotationally averaged collision cross section (CCS) value can be calibrated by measuring the drift time of proteins and protein complexes of known shape and structure.

Using this approach, we first examined vinculin. An IM–MS spectrum of vinculin shows a series of charge states separated in both m/z and drift time dimensions. The m/z projection of the IM-MS spectrum indicated 7 major charge states centered at 5,700 m/z (Figure 2A) corresponding in mass to monomeric vinculin. A calculated value of 6,531 ± 7 Å^2^ was obtained from the IM-MS measurements for the most intense charge state (+20) (Figure 2B). Theoretical CCS for human vinculin gave a value of 6,460 Å^2^, which fits well with the measured CCS value. Overall, these results suggest that in the absence of external perturbation, vinculin, in solution, exists entirely in the closed, head-to-tail packed, conformation.

**Figure 2.**
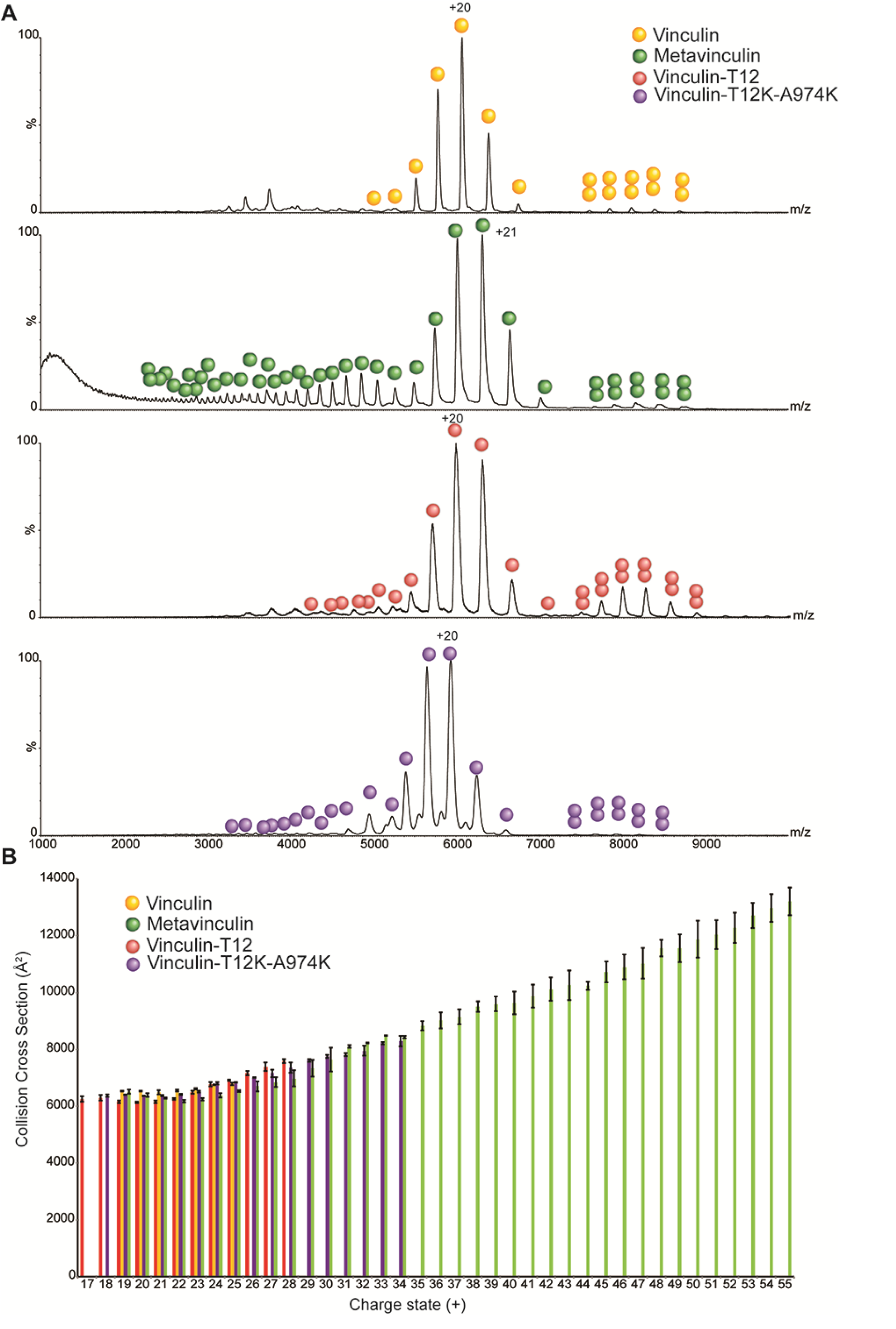
Vinculin variants exhibit altered conformational plasticity. Vinculin, metavinculin, vinculin-T12 and vinculin-T12-A974K were subjected to IM-MS analysis and their collision cross section was calculated. (A) Representative MS dimension projections of the four vinculin variants emphasizing the variability in charge state distribution. Both vinculin-T12 and to a larger extent metaviculin, display a wide distribution of charge states (29^+^–55^+^) compared to vinculin and T12. This heterogeneous, highly charged population indicates the existence of extended conformers. (B) Experimental CCS values. The high charge states of metavinculin display a significantly larger CCS value compared to the lower charge states (17^+^–28^+^) or to the CCS values obtained for vinculin and vinculin-T12. Error bars represent the standard deviation of 3 different wave heights.

Examination of metavinculin recordings indicated that it includes a considerably higher number of charge states, with a much broader distribution than that of vinculin (Figure 2). It was previously shown that the number of charges a protein can accommodate during the electrospray ionization process is correlated with its surface area (Hautreux et al., 2004; Kaltashov and Mohimen, 2005). Therefore, extended protein conformations, as well as partially or fully unfolded states, will give rise to higher charge states compared to those of the protein’s compact folded structure. Consequently, the larger number of charges observed for metavinculin suggests that in addition to the folded state, additional populations, with larger surface area, exist. This assumption was then validated by the CCS measurement. Metavinculin yielded a wide range of CCS values, which could be roughly divided into two groups. One population (charge states 19^+^–24^+^) displayed CCS values similar to vinculin, while the second population (charge states 25^+^–56^+^) exhibited much larger CCS values, up to 13,200 Å^2^, indicating the existence of a large population of molecules (in the order of ∼ 40%) with extended or unfolded states (Figure 2B). The conformational flexibility of metavinculin is likely to originate from its 68 amino acid insert residing between the tail and the hinge domains. Notably, this region exhibits no electron density in the crystal structure, suggesting that it is disordered. Taken together, these results suggest that the inserted segment of metavinculin modifies the dynamics of the head-to-tail lock, enabling metavinculin to form a more heterogeneous and extended set of conformations than vinculin.

We then set out to examine whether vinculin-T12 and vinculin-T12-A974K, in solution, are constitutively open, as predicted, or closed, as suggested by X-ray crystallography. The m/z projection of vinculin-T12 and vinculin-T12-A974K showed that they exhibit 11 and 16 charge states, respectively (Figure 2A). This charge state distribution is considerably narrower than that of metavinculin, however it is wider than that of vinculin, suggesting that vinculin-T12 and to a larger extent vinculin-T12-A974K, possess a somewhat greater structural heterogeneity than vinculin. The experimental CCS value for the highest charge state peak corresponded to 6,137 ± 29 Å^2^ for vinculin-T12 and 6,359 ± 9 Å^2^ for vinculin-T12-A974K. Thus, despite the elimination of three hydrogen bonds that anchor the head to the tail, these domains remain in contact in vinculin-T12 and vinculin-T12-A974K, in consistence with the crystal structure.

### Disruption of the head-to-tail interaction of vinculin is a two-step process

To explore the externally-induced conformational transition, associated with vinculin activation, we exposed the four vinculin variants to the Collision-Induced Unfolding (CIU) approach, which couples collisional molecular perturbation with ion mobility measurements (Hopper and Oldham, 2009; Zhong et al., 2014). This method elicits the conformational changes the ion undergoes as a function of the energy imparted. The collision voltage at which the transitions between conformations occur, the mode of the transition and the size of intermediates, generate a characteristic unfolding trajectory of the protein. Specifically, upon collision activation we expected that the head-to-tail constrains of the four vinculin forms will be resolved, yielding distinct CIU characteristics.

Considering that for low charge states the number of CIU transitions is correlated with the number of domains, indicating their uncoupling or unfolding (Zhong et al., 2014), we have reduced the number of charges on the proteins by utilizing triethylammonium acetate and have chosen to focus on the 17^+^ charge state, which for all vinculin forms corresponds to the closed compact head-to-tail conformation. Charge reduction also induces separation between protein peaks of similar mass, as their total mass is now divided by a smaller number of charges. Consequently, it was possible to simultaneously measure the CCS plots of a mixture of vinculin forms, while avoiding overlapping peaks and experimental variability. Towards this, experiments were performed in two series, one containing vinculin, metavinculin and vinculin-T12, and the other containing vinculin, metavinculin and vinculin-T12-A974K.

We first examined the unfolding properties of native vinculin. The CIU contour plot shown in Figure 3A, reflects the changes in the CCS values as a function of collision voltages. At low collision voltages, vinculin has an average CCS value of 6,266 ± 35 Å^2^, which is in agreement with the calculated CCS for the closed compact conformation of vinculin (6490 Å^2^) we therefore termed it the C (closed) state (Table 3). This configuration persists until reaching a collision voltage of 180 V, however, already at 140 V a transition occurs to a more extended, most likely “semi-open” conformation (state SO), giving rise to a CCS value of 6,970 ± 31 Å^2^. The transition of vinculin between states C and SO is a gradual process, whereby the increase in acceleration voltage leads to decreased levels of the compact form and a reciprocal increase in the SO state. An additional structural transition is observed at collision energies larger than 180 V, exhibiting a further increase in the CCS value, reaching 7,268 ± 15 Å^2^. This conformation persists when the maximal voltage limit is reached; we therefore termed it the “open structure” (state O). Overall two distinct and well defined ensembles of conformations are detected in addition to the most compact protein configuration. For comparison with the other vinculin forms, we use a simple (C, SO and O) nomenclature for these conformational transition families.

**Table 3:** Experimental collision cross measured from collision induced unfolding experiments. Values represent averages and standard deviations of 3 independent experiments in Å^2^.

| Protein\Conformation | C | SO | O |
| --- | --- | --- | --- |
| Vinculin | $6266 \pm 35$ | $6970 \pm 31$ | $7268 \pm 15$ |
| Metavinculin | $6281 \pm 32$ | $7056 \pm 30$ | $7363 \pm 56$ |
| Vinculin-T12 | $6108 \pm 38$ | $6847 \pm 28$ | $7173 \pm 125$ |
| Vinculin-T12-A974K | $6164 \pm 28$ | $6850 \pm 18$ | $7279 \pm 5$ |

**Figure 3.**
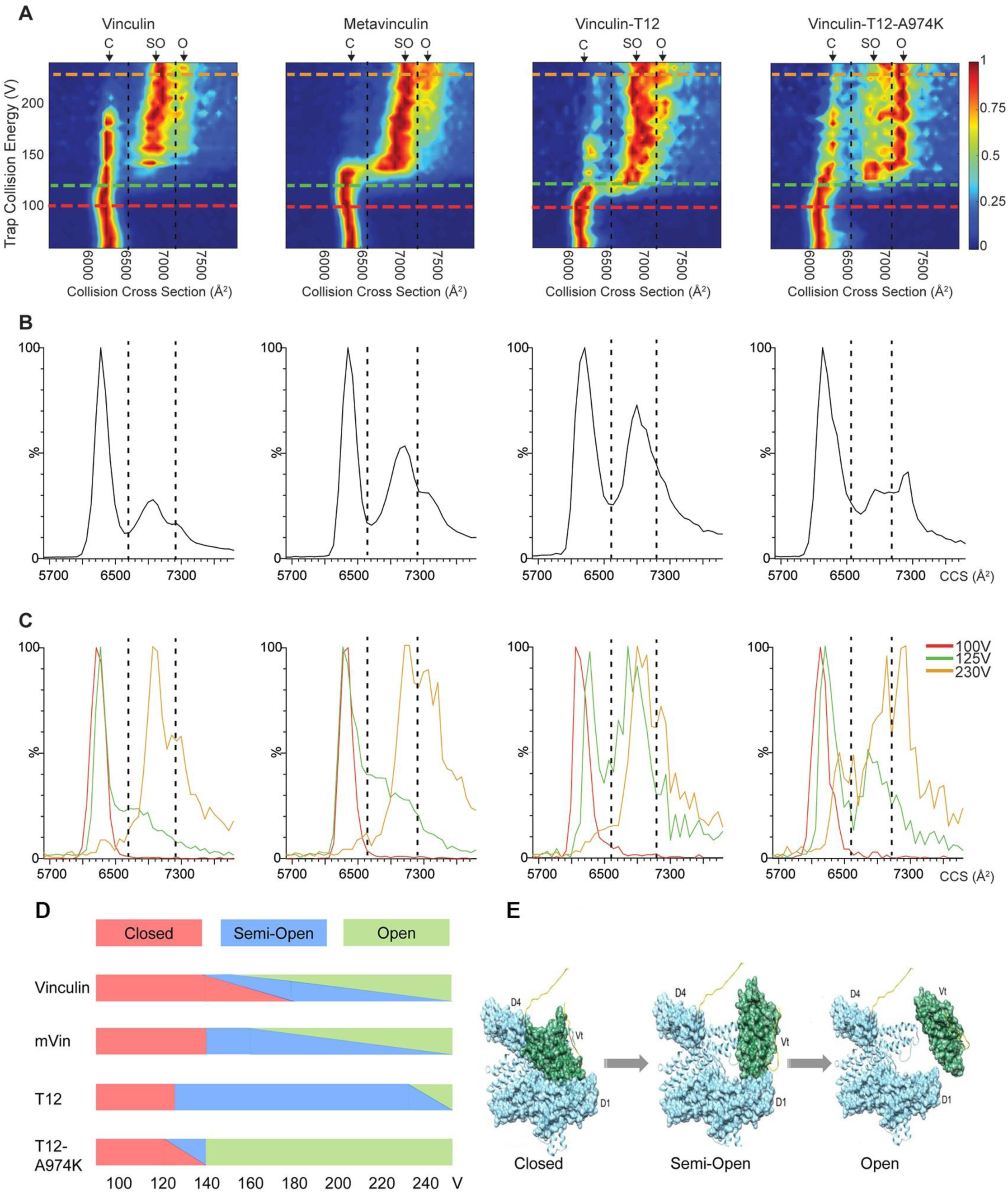
Collision induced conformational transitions in the four vinculin variants indicate the existence of a two-step opening mechanism. (A) CIU contour plots of the 17^+^ charge state of vinculin, metavinculin, vinculin-T12 and vinculin-T12-A974K, where CCS is plotted against collision voltage. Colors denote signal intensity as indicated. A similar unfolding pattern involving the two transition states from closed (C) to semi-open (SO) and consequently to open (O) conformations is observed for all proteins. Nevertheless, they differ in the energy threshold that is required for the transitions as well as in the range of voltages they dwell in each state. For clarity dashed lines were added. (B) Average CCS plots obtained for all measured collision voltages, reflecting the relative abundance of the three discrete states. (C) Superposition of CCS profiles obtained at collision energies of 100, 125 and 230 V emphasize the differences in the transitions state threshold of the various vinculin forms. The image is a representative experiment out of 4 repetitions. (D) A graphic representation of the data shown in panels (A) and (B). This illustration emphasizes the gradual transition of the C form of vinculin to SO. The transition of vinculin between the C to SO states is gradual, in contrast to the more abrupt shift of the other vinculin forms. Moreover, the transition of the two T12 forms occurs at a lower acceleration voltage, compared to vinculin and mVin. In addition, the apparent instability of the SO conformer of T12-A974K, compared to all other forms, and, in particular, to T12 is reflected. (E) Structural models of the proposed transitions between closed, semi-open and open conformations of the different vinculin variants.

The SO peak is clearly separate from the C state peak, suggesting that it represents a discrete state, distinct from the closed conformer. Therefore, in order to relate the measured collision cross sections to the structural changes that vinculin undergoes, we calculated the CCS for several in-silico models in which Vt of native vinculin was shifted away from Vh to a variety of locations. There was no one-to-one correspondence between the models and the calculated CCS values, namely, different models had similar CCS values. Yet, the calculated CCS increased as the distance between Vt and Vh increased and it reached 8,325 Å^2^ when Vt residue 875 was placed 125 Å away from D4 residue 836, a distance that approximately spans the length of an extended hinge. Figure S1 presents selected models that are fitted to the experimental CCS values (Table 3). In the C state, the head (Vh) domains D4 and D1 bind to the tail (Vt) domain. The SO state of vinculin, in which the interaction of Vt with either D4 or D1 is disrupted, is represented by two possible models, SO1 and SO2, respectively (Figure S1). The CCS calculations do not distinguish between the two options but the fewer and less energetic anchoring spots (see Table 2) at the significantly smaller interface of Vt-D4 compared to Vt-D1 strongly suggest that the Vt-D4 interface disrupts more readily than the Vt-D1 interface. Hence, the SO state likely corresponds to a conformation in which Vt is still in contact with D1 but not with D4. The CIU results indicate that C and SO conformers co-exist for a long range of collision voltages suggesting structural reversibility between the closed and semi-open conformations. The calculated CCS for an open vinculin, in which Vt is detached from Vh but does not move far away from it (Figure S1), is in correspondence with the O state. The CIU peak for the open state appears only after the closed conformer disappears and it merges with the SO state, suggesting continuous and reversible transition from semi-open to open conformers but not from closed to open conformers.

Inspection of the CIU plot of mVin, shows a similar pattern of two transition steps. Detailed analysis, however, reveals that unlike vinculin, the conformational transition from C to SO in mVin is abrupt; occurring at ∼140 V, whereby the C conformer of mVin disappears and the SO state simultaneously appears. This observation reflects different transition dynamics in mVin that can be attributed to the longer Vh-Vt linker. Thus, once the Vt-D4 interface is disrupted the longer linker hampers reversibility and affects the equilibrium between the SO and O conformers, explaining the more pronounced presence of the O state in mVin. The lack of reversibility in the C-to-SO transition in mVin is also in line with the accumulation of high levels of unfolded molecules in solution (Figure 2).

Interestingly, vinculin-T12 shifts between the compact C state and SO much earlier than vinculin, at the relatively mild conditions of 120 V. Since the T12 mutations at the Vt-D4 interface weaken this interaction, the observed early transition from the C to the SO state further supports our conclusion that SO represents a semi-open conformation in which Vt remains bound to D1 but detaches from D4. Hence, because of the weaker Vt-D4 binding, less energy is needed in order to induce the transition from closed to semi-open T12. Furthermore, the C of T12 state does not persist at higher collision voltages as for vinculin, probably because reversibility is hampered by the mutations. Interestingly, the C state of vinculin-T12-A974K persists longer than that of the T12 variant, but the SO state is detected at low voltage just like in T12, but is considerably less stable, and occurs for only a narrow range of collision voltages, before the next transition to the O state takes place. This higher stability of the C state in vinculin-T12-A974K (compared to vinculin-T12), is attributable to the hydrophobic contacts between the CH_2_ groups of K974 and D4 seen in the crystallographic structure. Stabilizing electrostatic interactions between K974 and D4 are not observed in the crystal structure of vinculin-T12-A974K, however, they can occur in solution or under the MS experimental conditions.

Taken together, these results show that all forms of vinculin exist in a closed conformation in the absence or at low level (<120V) of collision voltage (Figure 3D). Mutations that disrupt the head-to-tail binding (vinculin-T12 and vinculin-T12-A974K) can affect both the C-to-SO and SO-to-O transitions. The unexpected effect of the mVin insert indicates that the extended unstructured hinge region in mVin affects the C-to-SO transition. These results are in line with the crystal structures that show two-end binding of Vt to the D1 and D4 domains of Vh, forming two separate and apparently partially independent head-to-tail links. In accordance with this assumption, the CIU profiles indicate that the opening of the different vinculin variants is a two-step process, switching from closed to semi- and fully-open states (Figure 3E).

### The differential effects of vinculin and its variants on focal adhesion morphology and dynamics

Considering the fact that the four vinculin variants studied here are similar, in the sense that they all contain the same functional domains and all localize to FAs, yet they greatly differ in their head-to-tail lock strength, we examined the functionality of each of the four forms in live cells. To this end, we expressed each of the variants in two cellular systems, namely vinculin-null mouse embryo fibroblasts (MEF), where the transfected proteins are the only vinculin proteoforms present. In addition, we used HeLa cells, which also express an endogenous wild-type vinculin. In both cases, the transfected vinculin variants were fluorescently-tagged, enabling live cell monitoring of focal adhesion morphology and dynamics. Representative images are shown in Figure 4A and quantification of FA features is presented in Figure 4B-E. The analysis is based on quantification of approximately 1,000 FAs in each experimental group. As can be seen, the morphometric analysis revealed highly significant differences in FA area (Figure 4B and D) and length (Figure 4C and E), which were consistent in both the HeLa cells (Figure 4B and C) and the vinculin null MEFs (Figure 4D and E). In both cell types, vinculin expression correlated with enrichment of small (<1μm^2^) and short (<2μm) FAs, while T12 and, especially T12-A974K, being the largest and longest. The translocation of FAs in the different cells was variable, but consistently, cells containing vinculin were more dynamic than those expressing T12, T12-A974K and mVin (see supplementary movies 1 and 2).

**Figure 4.**
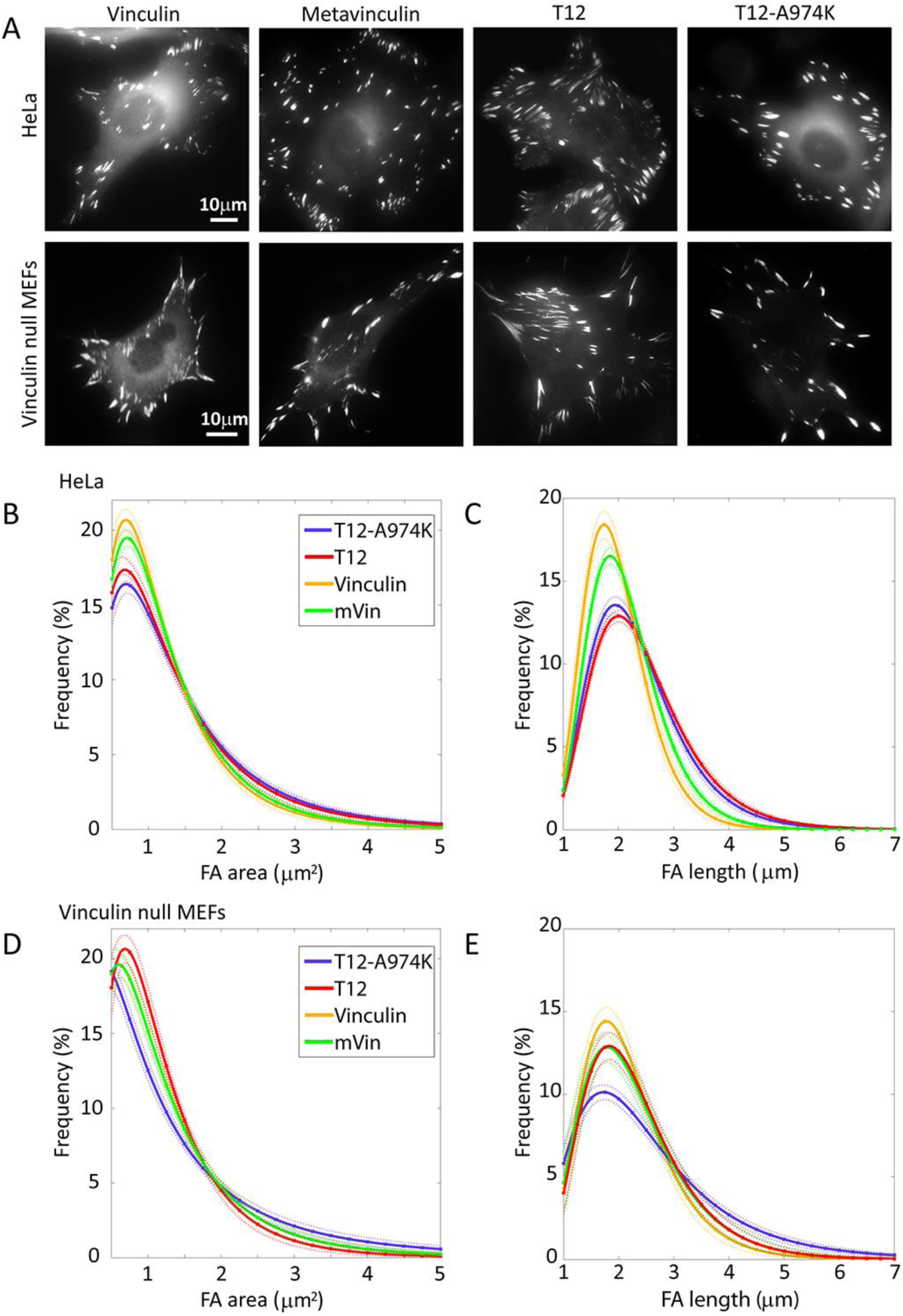
Vinculin variants have distinct effects on FA morphology. (A) The fluorescently-tagged vinculin variants localize to FAs. Scale-bar is 10 μm. (B-E) Quantification of focal adhesion properties of HeLa cells and vinculin null MEFs, expressing the four vinculin variants. Solid curves represent best fits to log-normal distribution profiles with 95% confidence bounds (dashed lines). (B) Distribution of focal adhesion area in HeLa cells. (C) Distribution of focal adhesion length in HeLa cells. (D) Distribution of focal adhesion area in vinculin null MEFs. (E) Distribution of focal adhesion length in vinculin null MEFs. These analyses are based on quantification of about 1,000 FAs for each experimental group.

### The differential effects of actomyosin contractility inhibition on the fate of FAs in cells expressing the different vinculin variants

To substantiate the relevance of the CIU data for the process that takes place in living cells, we have exposed vinculin null MEFs and HeLa cells, expressing fluorescently-tagged vinculin, mVin, T12 and T12-A974K to the Rho-kinase inhibitor Y27632, and monitored, by live cell fluorescence microscopy, the FA-associated fluorescence. The concentration of Y27632 used in these experiments was carefully calibrated, choosing relatively low levels of the inhibitor, which induce partial dissociation of FAs in WT vinculin-expressing cells. The low Y27632 concentration enabled us to record both stabilization and destabilization of FA in the presence of the different vinculin forms.

As shown in Figure 5 and in supplementary movies 1 and 2, vinculin expressing cells (both vinculin null MEFs and Hela cells) displayed a similar decline in FA fluorescence, with an initial slope of ∼1.5 %/min (in Hela cells) and 1.2 %/min (in vinculin null cells. Expression of T12 completely stabilized FAs in the vinculin null MEFs, and partially stabilized FAs in the HeLa cells, which express an endogenous WT vinculin (initial decline of fluorescence of 0.7%/min). Interestingly, mVin and T12-A974K, both increased the sensitivity of FAs, in both cells, to Y27632.

**Figure 5.**
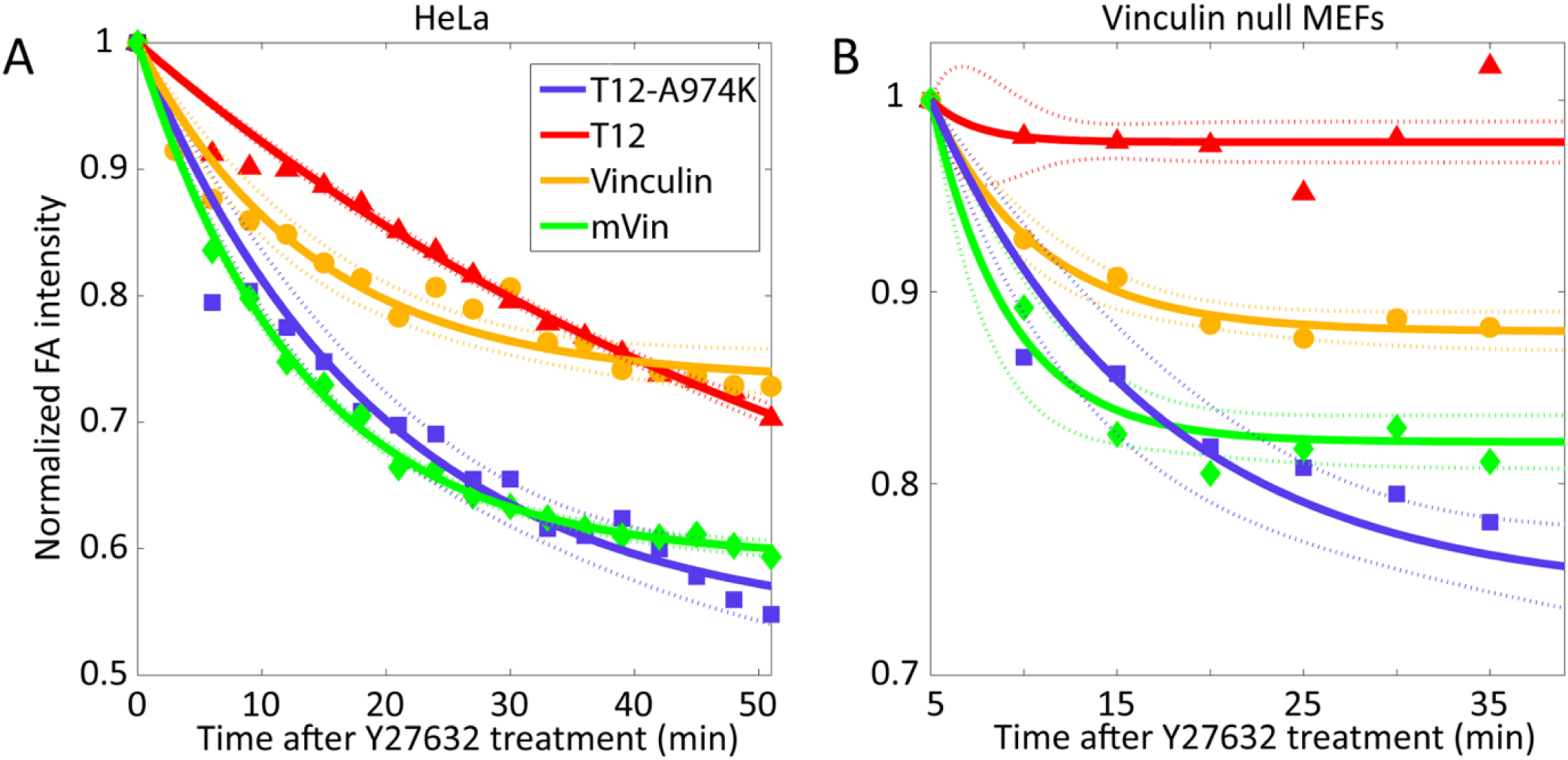
Inhibition of actomyosin contractility has distinct effects on FAs containing different vinculin variants. Time response of the different vinculin variants to Y27632 treatment in (A) HeLa cells and (B) vinculin null MEFs. Whereas T12 completely (vinculin null MEFs) or partially (HeLa cells) stabilized FAs, mVin and T12-A974K had the opposite effect of increasing FA instability. Solid curves are best fits to an exponential decay profile with 95% confidence bounds (dashed lines). Mean focal adhesion intensity was normalized with respect to its value just upon addition of 10μm Y27632 at time=0, and corrected for bleaching.

Taken together, the transition profiles of the four vinculin forms from closed conformation to semi-open to open bear a compelling correlation to FA structure, dynamics and mechanosensitivity. The two forms that most readily underwent transition from closed to semi-open conformation induces the largest FAs, while those forms that displayed conspicuous levels of an open conformation, like mVin (which displayed high levels of open molecules, even in the absence of external perturbation) and T12-A974K (which has a highly unstable SO form) produced FAs that were most sensitive to the stress blocker. Interestingly the “classic” T12, with its prominent levels of stable SO conformation, across a broad range of perturbation forces, supported the formation of large and force independent adhesion sites.

## Discussion

In this study we have explored the molecular mechanism underlying the force-dependent conformational extension of vinculin and its key role in the assembly of FAs. The widely accepted notion that vinculin is involved in mechanosensing at FAs is based primarily on structural and biochemical characterization of the protein. X-ray crystallography studies revealed a closed conformation of the molecule, in which the interaction of the “tail” domain with the “head” masks multiple binding sites to other adhesome molecules (Bakolitsa et al., 2004; Ziegler et al., 2006). The idea that the closed form of vinculin is “auto-inhibited”, and that pulling the tail domain away from the head might activate the molecule was supported by binding studies of isolated vinculin and, in particular, by studies conducted with the vinculin T12 mutant (Cohen et al., 2006). In T12, one of the head-binding sites on the tail was mutated resulting in a stably active form of vinculin, which increased the size and stability of FAs (Carisey et al., 2013; Humphries et al., 2007) and claimed to be constitutively open.

Our objective, in this work, was to monitor the conformational states of vinculin following the exposure to external perturbation. To determine the contribution of the key sites, which are expected to regulate the head-tail interactions, we compared vinculin with three variant molecules: metavinculin, which a naturally occurring alternatively-spliced form, present in muscle cells, in which the unstructured hinge region, inter-connecting the head and the tail is extended (Rangarajan et al., 2010). As well as two mutants which include modifications in the interaction site of the tail with the D4 domain of the head – one with four neutral alanine substitutions (the “classical” T12) and the other with a lysine residue, replacing alanine 974 in T12, and the aspartate in the original vinculin (T12-A974K). We chose this variant of T12, since we expected the replacement of the negatively charged aspartate with the positively charged lysine to further destabilize the head-tail interaction.

The first question, addressed by X-ray crystallography, was whether the T12-A974K is indeed a constitutively open molecule. Surprisingly, comparison of the new T12-A974K structure with the published structure of WT vinculin and of mVin indicated that despite the loss of 4 hydrogen bonds and the weakening of the remaining ones, the crystal structure clearly displayed a closed conformation. This can possibly be attributed to the strong binding of the head domain D1, which directs the other end of the rod-like Vt to bind to D4.

Being aware of the possibility that the crystal structure does not always represent the proteins’ dynamics in solution, we have subjected vinculin and the three variants to IM-MS analysis, which enables the derivation of experimental CCS values that are related to the surface area of a protein in solution (Bohrer et al., 2008; Wyttenbach et al., 2014). The IM-MS derived CCS values of vinculin, T12, T12-A974K and mVin indicated that these proteins exist in a closed conformation, supporting the X-ray crystallography results. T12-A974K and to a larger extent mVin, however, also possessed a population of conformers exhibiting larger CCS values, suggesting that the proteins exist, to some degree, in a partially unfolded conformation. In the case of mVin this is probably caused by its unstructured long insert positioned between the tail and the flexible hinge region. As suggested below, although the dominant form of the vinculin variants is the closed conformation, the extended conformations of T12-A974K and mVin harbor biological significance.

We next used the CIU approach to assess the differential structural properties of the vinculin forms, in response to acceleration energies (Zhong et al., 2014). These experiments revealed several unexpected features of the collision-induced conformational transitions of the four studied molecules. An important observation is that for vinculin and each of the variants, there were three discrete conformational states. The first corresponds to the closed (C) conformation, the second, to a semi-open (SO) state, and the third – to an open (O) state. Calculated CCS values, based on model structures, suggest that an O state can be reached, in which the head and tail domains are detached, while maintaining the folded tertiary structure of each domain intact (Figure 3E). Moreover, the distinct states appear to be stable species that exist for wide ranges of acceleration voltages, with only one exception, namely the semi-open state of T12-A974K, which readily proceeded to the open state (Figure 3D). Thus, a modular path of structural transitions exists between the C and O forms of the vinculin variants.

Comparing the four tested molecules, some interesting insights should be emphasized (see Figure 3D): (i) the C to SO transition appears to be tightly regulated by the tail-D4 interaction, hence, the two T12 mutants require lower acceleration voltage (120 volts) to undergo this transition in comparison to WT vinculin and metavinculin. (ii) The mode of transition of vinculin from the C state to the SO (between 140-180V), is gradual and possibly reversible. Unlike vinculin, the three vinculin variants undergo more abrupt transitions between the C and SO states. (iii) An intriguing observation made here is that the SO state in vinculin, mVin and T12 is relatively stable, while in T12-A974K the SO state is highly unstable. Thus, although the C to SO transition of T12 and T12-A974K is initiated at the same acceleration voltage, T12-A974K rapidly transforms to a stable O state.

Armed with this information concerning the conformational transitions of the different molecules, we have tested the differential effects of vinculin, mVin and the two T12 variants on FA size, dynamics and force dependence in vinculin null MEF and in HeLa cells. While the results obtained with the vinculin null cells provide more accurate information about the properties of the over-expressed variant, the comparison of the HeLa cells to the MEFs enabled us to determine whether mVin and the T12 mutants can induce their effects even in the presence of the endogenous vinculin.

Examination of the live cell movies generated from the two cell types, carrying vinculin and the three variant forms, provided a large amount of high quality data. Other features were examined too, but the variability between cells in the same sample was too high to draw significant conclusions. On the other hand, our results indicated that the size of FAs is clearly higher in cells expressing either T12 or T12-A974K, which undergo transition to the SO state under the milder conditions (Figure 3). These results suggest that the SO state is the active conformation of vinculin as the more easily triggered and abrupt transition of vinculin-T12 from C to SO is enough to dictate focal adhesion formation and stabilization.

Another interesting observation is that T12 expression in cells converts FAs to force independence, when expressed in vinculin null cells while T12-A974K exerts the opposite effect - namely it destabilizes the adhesions (Figure 5). This striking difference between the two molecules, which differ in only one amino acid, can be attributed to the apparent instability of the SO state in cells expressing T12-A974K. It is noteworthy that mVin also destabilizes FAs, which raises another attractive possibility, namely that the presence of an open conformer in the cytoplasm might induce FA disassembly.

Taken together, the results presented here support the following scenario concerning the mechanical activation of the two physiological forms. Vinculin, which is expressed in essentially all cells, and metavinculin, which is specifically co-expressed in muscle cells: Vinculin appears to have a very stable closed conformation, which can gradually undergo a transition to the active, semi-open state, driven by actomyosin contractility. This gradual “unlocking” of the closed conformation appears to depend on the force applied to the Vt-Vh(D4) interface, which can break the four hydrogen bonds present at this site (T974-N773, K975-E772 and bifurcated R978-E775). The elimination of these four bonds in the two T12 mutant molecules leads, not only to unlocking at lower activation level, but also to loss of the force-tunable activation process, and acquisition of a sharp transition from close to semi-open state. Interestingly, metavinculin started to undergo C-to-SO unlocking under the same level of mechanical perturbation as vinculin, but the process was abrupt, suggesting that the extra stretch of 68 amino acids at the hinge-tail interface affects the dynamics of the Vt-D4 interface. It is noteworthy that in solution, a significant proportion of metavinculin was already in the open state, which raises an intriguing question, whether the fully open vinculin or metavinculin can trigger or stabilize focal adhesions. We do not have direct information on the activity of the fully open vinculin and, in fact the results presented here support just the opposite, namely that the open forms (metavinculin and T12-A974K, with its highly unstable semi-open state) render them more sensitive to actomyosin inhibition.

## Author contributions

D.S.C., M.S. and B.G. planned the experiments and analyzed the data. D.S.C. performed the MS experiments. T.V. and A.L. performed the live-cell imaging experiments. T.V., A.L. and Z.K. analyzed the live-cell imaging data. M.E. performed computational analysis and modeling. B.M. and O.M. cloned, crystalized and solved the structure of vinculin-T12-A974K. B.M.J. cloned metavinculin and provided critical input on the manuscript. D.S.C., M.S. B.G and M.E. wrote the manuscript. Authors have no conflicting interests.

## Acknowledgments

We thank the Israel Structural Proteomics Center (http://www.weizmann.ac.il/ISPC) for cloning and purifying all the recombinant proteins used in this paper. We would also like to thank Brandon Ruotolo’s group for supplying the python scripts used to create the CIU plots presented in this paper, and Barbara Morgenstern for critical review of the manuscript. In addition, we would also like to thank Heidi Roschitzki-Voser for helping in the initial stages of the project. MS is grateful for the support of a Starting Grant from the European Research Council (ERC) (Horizon 2020)/ERC Grant Agreement no. 636752, and a Minerva Foundation grant, with funding from the Federal Ministry for Education and Research, Germany. BG would like to acknowledge support by the European Union Seventh Framework Program ERC Advanced Grant under grant agreement n ° 294852-SynAd and by the Israel Science Foundation, grant no. No. 3001/13. BG is the incumbent of the Erwin Neter Professorial Chair in Cell and Tumor Biology.

## Methods

### Cell culture and reagents

HeLa JW and vinculin-null MEFs were grown in DMEM (Gibco, Grand Island, New York) containing 10% FCS and 100 U/mL PenStrep (Biological Industries, Beit Haemek, Israel) at 37° C in a 5% CO2humidified atmosphere. ROCK inhibitor Y27632 (Sigma-Aldrich, St. Louis, MO) was used at a concentration of 10 μM.

### Overexpression of vinculin, metavinculin, vinculin-T12-A974K and vinculin-T12

HeLa and vinculin-null MEF cells were seeded onto optical glass bottom 24 well plates (MatTek, Ashland, MA) and then transfected with the different vinculin variants. HeLa cells were transfected using jetPEI (Polyplus-transfection, France), while vinculin-null MEF cells were transfected using either Lipofectamine 2000 or TurboFect (both Thermo Fisher Scientific, Waltham, MA). All transfections were done according to the manufacturers’ protocols. The vinculin variants were conjugated with GFP (except for vinculin-T12-A974K which was conjugated with mCherry) as described in (Zamir et al., 1999).

### Immunofluorescence microscopy and image analysis

Images and time lapse movies were acquired using a Deltavision Elite (GE Healthcare/ Applied Precision, USA) system mounted on an inverted IX71 microscope (Olympus, Japan) connected to a Photometrics CoolSNAP HQ2 camera (Photometrics, Tucson, AZ). The system was running SoftWorX 6.1.3. Pictures were taken with an Olympus UIS2 BFP1 60x oil PlanApoN objective with a numerical aperture of 1.42 (Olympus, Japan). Quantification of FA area, length and fluorescence intensity was performed using a custom code written in Matlab (MathWorks, Natick, MA) after correcting for fluorescence bleaching.

### Protein purification

The genes encoding WT Vinculin (2-1066), metavinculin (68 AA insertion following residue 915 of the wt sequence) and Vinculin-T12 (with the following mutations: D974A, K975A, R976A and R978A) were each cloned into pET28_TEVH. All three proteins were produced an purified in the same manner as follows: the plasmids were transformed into E. coli BL21(DE3) and a 5L culture was grown in LB medium at 37°C until mid-log phase. Protein expression was induced by the addition of 0.2 mM IPTG and allowed to continue growing at 15°C ON. After harvesting the cells by centrifugation, the cell pellet was lysed using a cell disrupter (Constant Systems) in 100ml buffer containing: 50mM Tris pH=8, 0.5 M NaCl, 20mM Imidazole, protease inhibitor cocktail set III (Calbiochem), Dnase, 1 mM PMSF and Lysozyme. The insoluble material was removed by centrifugation at 26,000 g for 30 min. The supernatant was filtered and loaded onto a Ni Column (HiTrap_chelating_HP, GE) equilibrated with lysis buffer. The protein was eluted in one step with the same buffer containing 0.5M Imidazole. Fractions containing Vinculin were loaded onto a size exclusion column (Hiload_26/60_Superdex200, GE), equilibrated with 50mM Tris pH 8 and 50mM NaCl. The pooled fractions containing Vinculin were injected into an anion exchange column (Tricorn Q 10/100 GL, GE) equilibrated with 50mM Tris pH=8. Pure Vinculin was eluted with a linear gradient to 1 M NaCl. The pure protein was flash frozen in aliquots using liquid nitrogen and kept at -80o C. For purification of vinculin-T12-A974K, Cultured Sf9 insect cells at a density of 1.5 to 2.5 mio/ml were infected with P1 virus and incubated for ∼27 hours at 27°C. Expressing cells (green color) were harvested and lysed in 20 mM Tris-HCl (pH = 7.5), 0.4 M NaCl, 1 mM DTT, 0.01% Triton, PMSF, and PI cocktail. The protein was purified over a nickel affinity column and eluted in presence of 3C protease. Vinculin was further purified in 20 mM Tris-HCl (pH = 7.5), 0.1 M NaCl and 1 mM DTT using a size exclusion column (Superdex 200 Increase 10/300 GL). Protein concentration was determined, aliquoted and stored at -80°C.

### Crystallization

Purified vinculin T12-A974K (∼6mg/ml) was crystallized using the vapour diffusion, sitting drop method at 4°C. First crystals emerged from initial screens and were used for microseeding, yielding new crystals gained against a reservoir solution consisting of 15% (w/v) polyethylene glycol (PEG) 4K, 0.2 M calcium acetate, 0.1 M Tris-HOAc (pH 8.5), and 15% ethylene glycol (EG) for cryoprotection. A total of 70 crystals were fished and flash frozen in liquid propane.

### X-ray diffraction data collection

Data sets were collected on frozen crystals on the Xo6DA (PXIII) beamline at the Swiss Light Source of the Paul Scherrer Institute on a PILATUS 2M detector (Dectris). Crystals diffracted to a resolution of ∼2.9 and belonged to the trigonal space group P3121 (a = b = 97.81, c = 233.77, α = β = 90, γ = 120).

### Structure determination and refinement

The data were indexed, integrated and scaled with XDS. Molecular replacement was done with Phaser using a vinculin WT (PDB code 1TR2) search model. Model building was performed with the program Coot. Initial refinement was done with Refmac and finalized with Phenix. Rigid body refinement cycles with data from 20.0 to 3.0 Angstrom were alternated with manual model building and the R/Rfree values were monitored throughout this process. The final model shows R/Rfree values of 25% and 30.6%, good geometry and almost no residues in disallowed regions of the Ramachandran plot.

### Computational anchoring spots mapping and molecular modeling

We used a modified version of ANCHORSMAP (Ben-Shimon and Eisenstein, 2010), which includes threonine and proline anchoring spots mapping, to detect anchoring spots on the surface of native vinculin. Anchoring spots are energetically favorable binding locations of single amino acid side chains. Each anchoring spot consists of a binding cavity on the surface of a protein, bound to an amino acid side chain. The procedure detects small cavities on the surface of a protein, scatters thousands of amino acid probes near the cavities and determines optimal probe positions. The binding ΔG of the optimally posed probes are calculated employing a semi-empirical scoring function that includes van der Waals and solvation energy terms and an electrostatic energy term corrected for the dielectric shielding exerted by an approaching protein. In this study the positions of low ΔG anchors (ΔG≤-3 Kcal/mol) were compared to the X-ray structure of human vinculin (PDB code 1TR2), searching for anchors that closely corresponded to observed binding sites. The UCSF-Chimera software (Pettersen et al., 2004) was used for structure analyses, anchoring spots visualization, preparation of the semi-open and open models of vinculin and preparation of figures 1 and 4.

### Ion-mobility mass spectrometry

All ion mobility measurements were recorded on a Synapt G1 HDMS system (Waters Corp., UK) with a traveling-wave ion mobility device as described in detail elsewhere (Michaelevski et al., 2010). Proteins were first buffer exchanged into 200 mM 19rimethy acetate, pH=7.6 (Sigma) using a Biospin 6 column (Bio-Rad) and their concentration was adjusted to a final concentration of 10 μM. Typically, protein aliquots of 2-3 μl were injected via a gold coated borosilicate capillary via a nano-ESI ion source. Acquisition parameters were as following-capillary voltage-1.25 kV, source temperature-25°C, sampling cone-17 V, extraction cone-1.1 V, trap and transfer collision energy-5 V. For IMS wave velocity of 250 m/s and wave heights of 8-10 V were used. IMS calibration for conversion of drift times to collision cross section values was done as previously described (Bush et al., 2010). Experiment was repeated three times. Data presented is from a representative experiment and is an average of three wave heights-8,9,10 V.

### Collision induced unfolding (CIU) assays

Collision induced unfolding experiments were performed on a Synapt G1 HDMS system (Waters Corp., UK). Initially, proteins were buffer exchanged into 200 mM ammonium acetate pH=7.6 supplemented with 40 mM trimethylamine (TEA). The latter was added in order to induce charge reduction and avoid a large number of overlapping peaks. Experiments were done in 2 series combining three proteins at once to reduce experimental variability. The first series included vinculin, metavinculin and vinculin-T12, while the other included vinculin, metavinculin and vinculin-T12A974K. Collision induced unfolding was performed as previously described (Laganowsky et al., 2014; Zhong et al., 2014). In brief, the proteins were activated by increasing the trap collision voltage from 50 to 240V by 5 V increments. To compensate for slow traveling times due to reduced charge, IM-MS acquisition parameters were set as follows-wave velocity of 250 m/s, variable wave height between 0 to 30V with a ramp time of 10%. IM-MS calibration for conversion of drift times to collision cross section values was done as previously described (Bush et al., 2010) using the instrument parameters presented above.

### Theoretical Cross-Section Calculations

Theoretical collision cross section was calculated using the CCScalc function of Driftscope 2.5 (Waters).

